# Biological machine learning combined with bacterial population genomics reveals common and rare allelic variants of genes to cause disease

**DOI:** 10.1101/739540

**Authors:** DJ Darwin R. Bandoy, Bart C. Weimer

## Abstract

Highly dimensional data generated from bacterial whole genome sequencing is providing unprecedented scale of information that requires appropriate statistical frameworks of analysis to infer biological function from bacterial genomic populations. Application of genome wide association study (GWAS) methods is an emerging approach with bacterial population genomics that yields a list of genes associated with a phenotype with an undefined importance among the candidates in the list. Here, we validate the combination of GWAS, machine learning, and pathogenic bacterial population genomics as a novel scheme to identify SNPs and rank allelic variants to determine associations for accurate estimation of disease phenotype. This approach parsed a dataset of 1.2 million SNPs that resulted in a ranked importance of associated alleles of *Campylobacter jejuni porA* using multiple spatial locations over a 30-year period. We validated this approach using previously proven laboratory experimental alleles from an *in vivo* guinea pig abortion model. This approach, termed BioML, defined intestinal and extraintestinal groups that have differential allelic variants that cause abortion. Divergent variants containing indels that defeated gene callers were rescued using biological context and knowledge that resulted in defining rare and divergent variants that were maintained in the population over two continents and 30 years. This study defines the capability of machine learning coupled to GWAS and population genomics to simultaneously identify and rank alleles to define their role in abortion, and more broadly infectious disease.

## Main

Comparative microbial genomics has emerged from pangenome comparisons that are exclusively tied to reference genomes that define the perspective of change to a core and flexible genome perspective lacking a firm confirmation of which genes are linked to disease^1^. An alternative approach to this perspective is use of genome wide association (GWAS) methods that are common in mammalian genomics in an effort to refine the estimates of specific genes of interest. A limitation of GWAS is that it sequentially examines single loci that prevents simultaneous analysis of different allelic variants that can be interacting at different levels and population distribution between strain differentation^2^. This is a severe limitation in bacterial genomics, especially as population genomics is now possible in bacteria at a scale that allows examination of non-linear evolutionary rates of each gene and all of the alleles found in very large populations that create big data analytical problems. A compounding limitation is the lack of appropriate statistical models that underpin this approach in bacteria since it is unknown when the populations are normally distributed or evolving in a non-linear progression. As with all large data sets, multiple comparisons require Bonferroni correction to adjust the *p*-value based on a new scale as compared to gene expression but it is on a scale that is beyond that contemplated for gene expression variation (Table 1)^3^. Further, the assumption that each gene or allele is independent is conceptually flawed; and hence, alternative analyses that are biological and statistically compatible needs to be defined.

**Table 1.** Exemplar comparison of statistical metrics of GWAS versus machine learning metrics. Allelic variant association with phenotype using XGboost. An allele can be very large, ~8,000 for porA for a pairwise comparison. Using a population of this gene from 200 genomes created a population variation of 1.2 million variants that can be ranked with an estimation of importance to association with the disease phenotype, abortion in this case.

|  | GWAS statistical metrics |  | Machine Learning coupled to GWAS metrics |  |
| --- | --- | --- | --- | --- |
| Allele | GWAS p-value | Bonferonni Corrected p-value | Candidate Ranking | Feature Importance |
| $X_1$ | 0.001 | $8.3 \times 10^{-10}$ | 1 | 80 |
| $X_2$ | 0.001 | $8.3 \times 10^{-10}$ | 2 | 75 |
| $X_3$ | 0.001 | $8.3 \times 10^{-10}$ | 3 | 70 |
| $X_n$ | 0.001 | $8.3 \times 10^{-10}$ | $\text{Rank}_n$ | $\text{Importance}_n$ |

Coupling GWAS, population microbial genomics, and machine learning is poised to be a robust alternative to classical GWAS or pangenome comparison to simultaneously discover changes in microbial genomes, and genes, that span the scale of genome plasticity to alleles of a single gene. Moreover, this combination (coined BioML) will produce a statistically underpinned comparative ranking of the most important factors that are not obvious from GWAS alone. These advantages combined with downstream inspection of the prioritized rank further powers discovery to bring biologically insightful observations and solutions, especially when large genome populations are used in the analysis, from very divergent populations of alleles that are missed when gene calling is too divergent.

An analytical strength for use of machine learning in microbiology is the ability to define functional relationship from population scale genomes or genes without a priori definition of the underlying mechanism of change or specific phenotype limitations^4^. This distinctive advantage makes machine learning superior to classical statistical tests for prokaryotic systems that are highly variable, particularly bacteria wherein explanatory variables are not linearly correlated, features are dependent due to genome linkage, varying evolutionary rates of between genes, and assumptions of normal distribution are violated in part due to varying selection conditions^2,5^. These biological conditions and parameters are incompatible with the assumptions of linear or correlative statistics, which is compounded with data reduction methods that provide a very small snapshot of the genome variation that yield associations that have low predictive value.

In this study, we verified the concept of coupling GWAS with machine learning and population bacterial genomics (Figure 1) in a use case to test the hypothesis that a specific gene (*porA*) is linked to extraintestinal location and further is causative in abortion^6–8^ in a ranked order that is biologically meaningful. This was done using a wet lab validated data set containing 100 genomes^6–8^ using extreme gradient boosting (XGboost), which was used in biological applications previously^9^. XGboost can identify genetic variants in human GWAS as demonstrated in a Finnish study that integrated complex nonlinear interactions of SNPs^10^. The ability to interrogate the predictive features enables whiteboxing the parameters, which is emerging as a tool for deriving mechanistic function in biology^11^. XGboost implements adaptive optimization within the functional space by iteration of the weak learners into strong learners represented by decision trees where each new decision tree is generated by factoring the residual generated from the difference from observed to the predicted feature (Figure 2; Supplemental Table 1).

**Figure 1.**
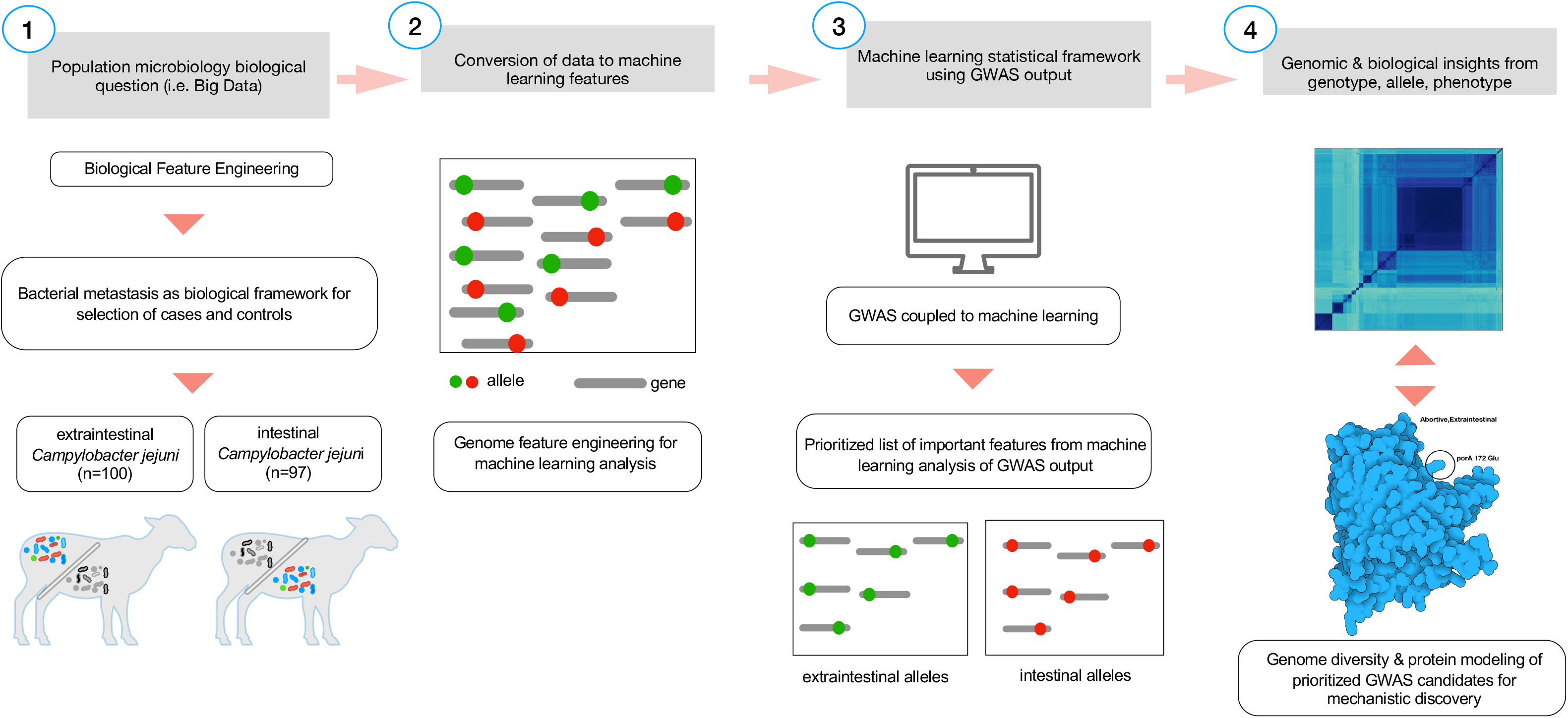
Biological feature engineering of genomic data for machine learning analysis. A critical step in feature engineering is selection of the appropriate comparison groups to enable classification of alleles that are related to the specific phenotype of interest (i.e. intestinal (controls; diarrheal; n=108) and extraintestinal (cases; abortive; n=85) (Step 1). Population-wide allelic variants (red dot = intestinal, green dot = extraintestinal) that result from variant calling (Step 2) and are used as the input features for machine learning analysis (Step 3). The predicted model generated from the machine learning analysis is inspected for the most predictive features using biological context, input, and protein modelling (Step 4) that represents a nonsynonymous mutation from the genomic the population of allelic variants (n=193).

**Figure 2.**
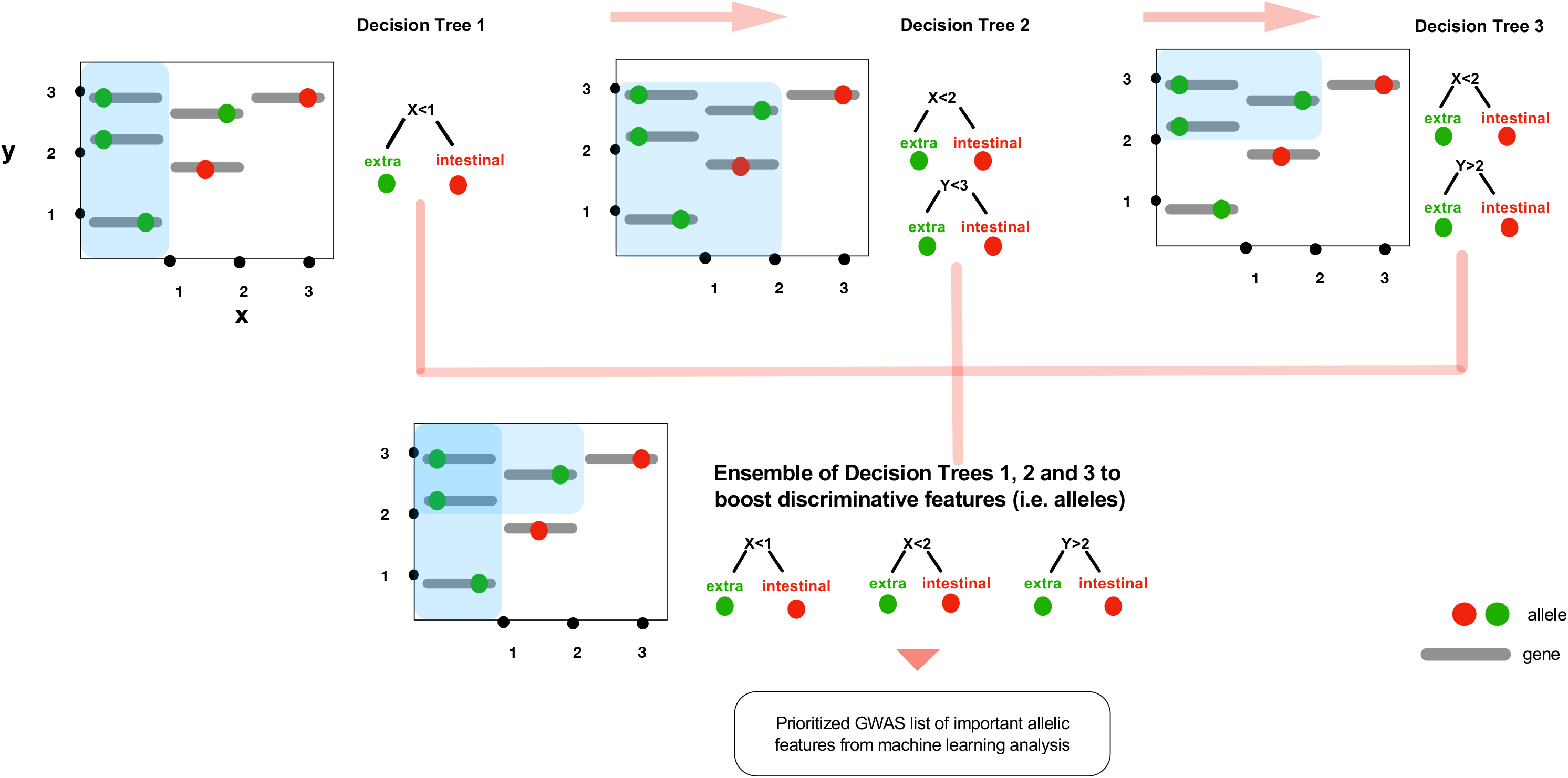
The conceptual framework diagram depicting machine learning in bacterial genome wide association using extreme gradient boosting (XGboost). Boosting is a technique of combining a set of weak classifiers or decision trees to increase prediction accuracy. Red dots represent an allelic variant, each grey bar represents a unique allele. Individual decision trees (1, 2, 3) fail to fully capture the allelic variants associated with the phenotype (e.g. extraintestinal abortion), but by combining the trees together results in a process called as boosting increases the discriminative power.

This study used a previously validated wet lab data set with a tetracycline resistant strain of *Campylobacter jejuni* causing abortion in sheep^6–8^. Their studies used a pairwise genome comparison to identify 8,000 SNP difference between a reference genome and abortive strain and utilized transformed genomes to identify specific allelic variants driving abortion. We utilized those 85 genomes that span 30 years and multiple locations as a reference set of cases and 108 control genomes of intestinal, diarrheal isolates. This approach allows exploration of bacterial population genomic space by linking different phenotypes to the genome variation among the isolates (Figure 1). Biological feature engineering of this collection of genomes identified 1.2 million SNPs, which is not tractable using in vivo infection studies to determine the roll of all SNPs. To examine this scale problem, we hypothesized that genomic changes evolved in gastrointestinal *C. jejuni* resulting in an abortive phenotype; hence, moving from the intestine to other tissues – in this case the placenta resulting in abortion. Applying our approach (BioML) to a population of gastrointestinal, diarrheal *C. jejuni* versus extraintestinal, abortive phenotypes produced a prioritized allelic difference in a ranked order of importance to the phenotype (i.e. abortion) (Supplementary Table 1).

BioML identified 14 *porA* loci as the most important alleles ranging from 89 to 59 relative importance out of the 1.2 million SNPs (Supplemental Table 1). These ranked loci were compared by body location (Figure 3), which further clarified the location of these SNPs in a Tetris plot that simultaneously presented the ranked associated allelic variants within the phenotype of interest as detected with BioML as well as the non-associated alleles. By presenting the cases and control simultaneously within the y-axis, capturing insight is easily observed and areas for further investigation can be prioritized with visual inspection combined with biological knowledge. An added feature of the Tetris plot, which is lacking in Manhattan plots, is the ability to detect rare variants that are not captured by gene calling, machine learning alone, or classical statistical testing. Regions within cases expressing different allelic patterns were further explored for each genome and implications in biological features important in the disease. Additionally, protein structures were modeled to examine the changes in protein configuration initially yielded three distinct groups (Figure 3). These alleles were directly compared to those validated *in vivo* and found to be linked to specific protein loops within alleles verified previously^6–8^ – BioML found each of those to be biologically important for abortion.

**Figure 3.**
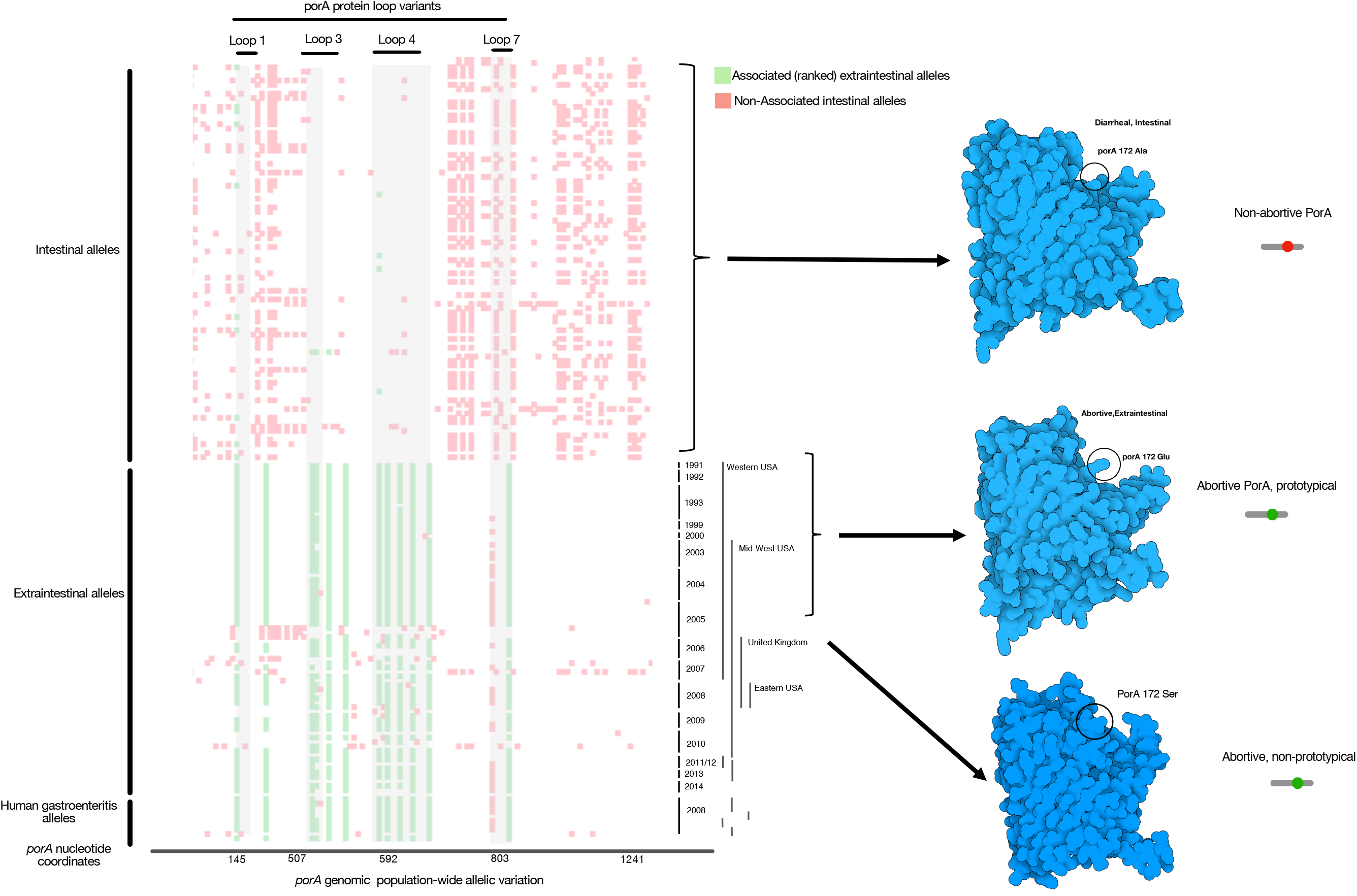
Comparative plot of SNP loci along the *proA* gene in all genomes. We termed this a Tetris plot as an alternative visualization of genome wide association hits because they are ranked and display only the loci that vary to produce a nonsynonymous mutation. The y-axis contains individual genomes from the cases and the controls, while the x-axis contains the GWAS SNP loci (green), the non-disease associated SNPs (red), open space (white) are loci that are identical in the gene sequence. Temporal and geographic metadata on the right side of the Tetris plot provides context for mutational enrichment over 30 years and multiple distant locations in North America and the UK. The enriched SNP variation produced different protein structures (far right in blue) as the corresponding protein model by location within the animal by SNP. Protein structural features corresponding to the ranked GWAS variants are annotated on top and below the plot are the nucleotide coordinates. Rare variants (homology <75%) was not included by the variant caller in this visualization but manual inspection provided a method to find these variants.

Further we located each of the top ranked alleles loops 1, 3, 4, 7 as enriched selection loci, again verifying previous wet lab observations^6–8^. Tetris plot derived variants that were not 100% identical with >75% protein homology, were designated as nonprototypical variants because the sequence variation was high enough to change protein structures. In a limited set of alleles, the *porA* allele was so divergent that they were not variant called but were recovered with manual curation for this study. Recovery of these genes that were not initially identified created a third group of rare variant alleles that also caused abortion (Figure 4; protein homology <75 %). All of the variants were mapped to the whole genome phylogeny diversity as well rare variants that were not variant called by the reference genome (Figure 4). Prototypical allelic variants clustered in the largest genomic group of abortive isolates, as did some of the nonprototypical *porA* variants. However, there was significant genome variation and contained the two groups that caused abortion. Rare *porA* variants were distributed within different genomic groups as well as over a 15-year span between North America and the UK. The extensive allelic versions of *porA*, as well as the different genotypes, suggests that a genome surveillance system based on SNPs would be unsuccessful to link these genomes to a disease. In combination, these observations indicate that BioML produced a ranked list of biologically important alleles that were validated with those that were previously shown to be causal in abortion for the exact SNP and the protein loop location. Together, these observations verified that BioML was capable of accurately identifying the exact SNPs in *porA* that cause abortion.

**Figure 4.**
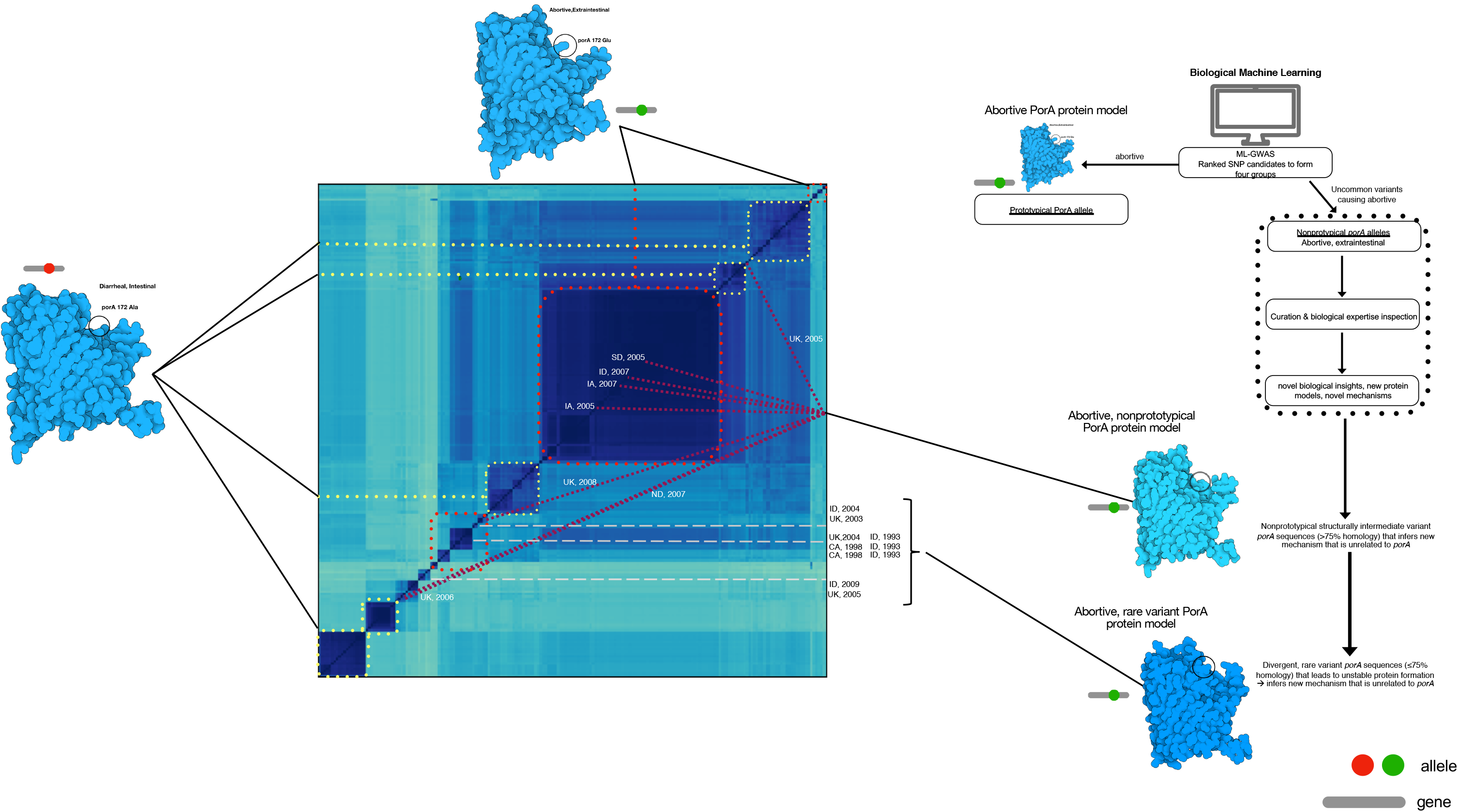
Whole genome distance matrix using minhash depicting an all against all comparison of genome diversity for all isolates used in this study overlaid with the porA variant associated with body location and disease phenotype. Genotypes and *porA* variants are connected in this depiction to examine the association between intestinal/diarrheal location (yellow dot boxes), prototypical extraintestinal/abortive (red dot boxes), non-prototypical *porA* variants in extraintestinal/abortive (maroon lines), and rare *porA* variants in extraintestinal/abortive (grey dashed lines) were co-located to their respective genomes in the genotype map. For the non-prototypical variants, the year and location of isolation was included to depict the variation over time and space in the maintenance of a minority population of *proA* variants of extraintestinal abortive *Campylobacter jejuni*. The diagram to the right depicts the process used for this analysis.

Since each BioML allele was validated for accuracy to wet lab results correctly, we further examined the protein changes from the ranked alleles (Figure 5). The first six top ranked alleles changed the amino acids for each *porA* sequence, but each protein sequenced varied across all PorA models. However, lysine_189_ was conserved across the extraintestinal variants and Asn was found in the intestinal alleles. Lysine mutation changes are the most impactful in membrane pore structure and are one of the tenets of membrane topology as positive inside rule^12,13^. Positive inside rule describes the observation across membrane pores that positively charged amino acids are found within the cytoplasm and negatively charged amino acids are in the extracellular domain. Membrane topology can radically change from being oriented inside the membrane (exposed to the periplasm in this case) to outside the membrane with a single lysine mutation. Within the adjacent protein structure, lysine snorkeling effectively minimizes the nonpolar chain component by burying in the hydrophobic domain and at the same time expose the polar component to the aqueous domain is another single amino acid change which alters the topology of the membrane domain^14^. Bacterial membrane pore flipping could be a potential mechanism to avoid recognition by the immune system and enhancement of ion transport. While the counterpart position is buried in a deeper position due to insertional mutation in rare variants, the inserted amino acids contain lysine at new position 197. Additionally, insertions in the rare variants reduce the homology to < 75 % lead to more extensive protein structural changes that change the PorA arrangement in the membrane while still able to cause abortion. This situation is troublesome for traditional approaches but BioML effectively identified this situation.

**Figure 5.**
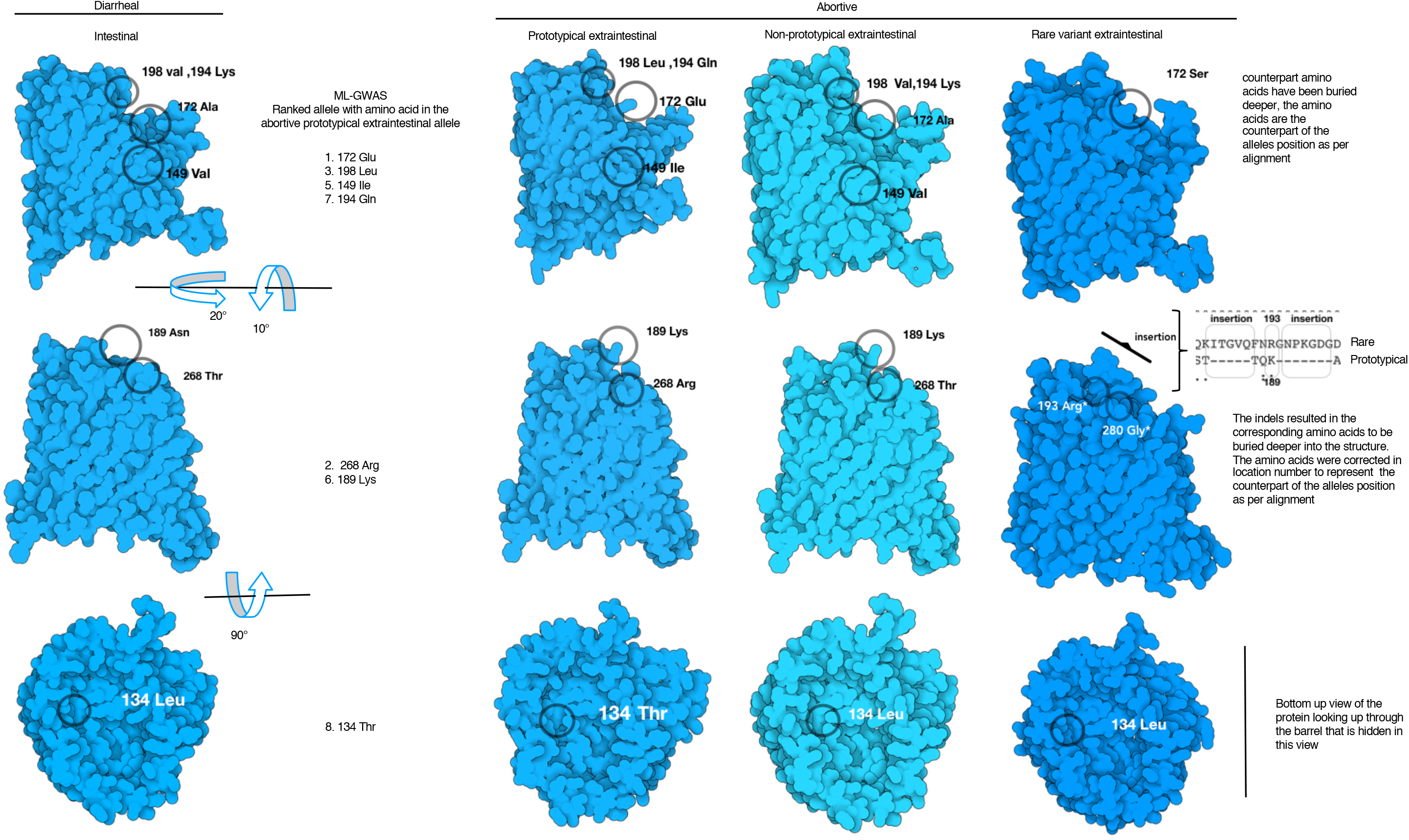
Protein models of the four groups of *porA* allelic variants that change the protein model structure relative to the isolate location in the host and the disease outcome. The amino acids corresponding with the BioML top ranked alleles are labelled in the common variant of *porA*, while the rest show the substituted amino acid in their respective position.

This study utilized a combination of GWAS, population bacterial genomics, and machine learning to identify and rank allelic variants that correspond to biologically validated alleles of porA to cause abortion. BioML results were further supported by the longitudinal and spatial conservation of *porA* coupled to protein substitutions that led to biologically relevant changes in the structure to change activity. A Tetris plot visualization provided an avenue to discover divergent and rare variants that provided further insight with protein modelling that uncovered protein substitutions resulting in localization changes that affect activity and isolation localization in the host. Together these results demonstrate and validate a novel method, termed BioML, to discover biological mechanisms using population bacterial genomics. This approach provides an avenue to leverage the massive amount of bacterial genomic sequences to uncover new mechanisms of disease with potential to provide therapeutic approaches.

## Acknowledgement

DDB is grateful for the funding provided by the Philippine California Advanced Research Institute and University of the Philippines Enhanced Creative Work and Research Grant to fund his PhD program.

## List of Supplemental Tables and Figures

Supplemental Table 1. Ranked allelic variants using BioML

Supplemental Table 2. Metadata for extraintestinal *Campylobacter jejuni*

Supplemental Table 3. Metadata for intestinal *Campylobacter jejuni*

Supplemental Table 4. Confusion matrix and derived model metrics for the XGboost model with extraintestinal *Campylobacter jejuni*. TP= True positive, FN= False Negative, FP= False Positive, FN= False Negative.

## Methods

### Biological feature engineering

Biological feature engineering entails selection of pertinent controls and cases for BioML analysis. The genomes between gastrointestinal and extraintestinal abortive isolates. *C. jejuni* controls were downloaded from Patric 3.5.28 (https://www.patricbrc.org/), June 1, 2019 (Supplemental Table 2). Abortive extraintestinal genomes *of C. jejuni* were obtained from the Sequence Read Archive (SRA; Supplemental Table 3)^8^. Fastq files were assembled using Shovill 1.0.4 (https://github.com/tseemann/shovill). Assembled files were annotated with Prokka (version 1.13.3)^15^. Variant calling was done with the reference sequence *C. jejuni* NTC11168 with Snippy 4.3.5 (https://github.com/tseemann/snippy) as previously described^16^.

### Gradient tree boosting as GWAS framework

GWAS variants generated from the biological feature engineering step were used as input for XGboost. The source code for implementing gradient tree boosting is available at https://xgboost.readthedocs.io/. Confusion matrix were generated and used to assess the performance of the model (Supplemental Table 4). The relative importance of the predictive model was used as the GWAS hits.

### Tetris plot

Classical GWAS hits are displayed as the negative logarithm of the p-value in Manhattan plots, hence we formulated a novel visualization of the ranked alleles generated by the machine learning model to highlight the difference between approaches - we call this GWAS hit visualization a Tetris plot. We color coded the relative importance values of the associated alleles derived from the XGboost (green being associated and red being non-associated). The source genome is plotted on the y-axis and genomic coordinates on the x-axis overlaid with GWAS hits presence or absence matrix.

### Population wide whole genome phylogeny

The genome distance metric was calculated using genome wide k-mer signatures to generate the population-wide phylogeny with a k-mer size of 31 scaled to 1000 with Sourmash^17^. The resulting genome wide k-mer distance was visualized as an all-against-all heatmap^17^.

### Protein Modelling

Assembled genomes were annotated using Prokka (V1.13.3) and PorA protein sequences were extracted for protein modelling using Swiss Model^18,19^. The most homologous protein was used as template for protein modelling. Illustrate (https://ccsb.scripps.edu/illustrate/) was used to generate the protein visualization of the predictive alleles. Ranked BioML alleles identified by visual inspection of the Tetris plot, via the ranked variable importance were used to inspect the protein structures.

