## Supplementary material for "Biological machine learning combined with bacterial population genomics reveals common and rare allelic variants of genes to cause disease": supp table 1

| allele | relative importance | scaled importance |
| --- | --- | --- |
| porA_513/1275 | 89.39 | 0.9998 |
| porA_803/1275 | 86.02 | 0.9621 |
| porA_592/1275 | 84.53 | 0.9455 |
| porA_548/1275 | 83.56 | 0.9346 |
| porA_445/1275 | 83.19 | 0.9304 |
| proS_1373/1707 | 83.12 | 0.9297 |
| porA_562/1275 | 82.20 | 0.9194 |
| porA_580/1275 | 81.20 | 0.9082 |
| porA_400/1275 | 81.16 | 0.9077 |
| porA_250/1275 | 79.46 | 0.8887 |
| porA_459/1275 | 79.24 | 0.8863 |
| ubiE_2_255/774 | 78.06 | 0.8731 |
| kpsM_157/783 | 77.53 | 0.8672 |
| hemL_762/1275 | 77.09 | 0.8623 |
| proS_1512/1707 | 76.92 | 0.8603 |
| sdhA_1599/1836 | 76.42 | 0.8548 |
| ackA_1114/1191 | 76.41 | 0.8547 |
| kpsD_1065/1659 | 76.37 | 0.8542 |
| porA_411/1275 | 75.90 | 0.8489 |
| tatC_94/738 | 75.65 | 0.8461 |
| proS_67/1707 | 75.64 | 0.846 |
| nrdA_315/2370 | 75.14 | 0.8405 |
| kpsD_55/1659 | 75.04 | 0.8393 |
| ubiD_1006/1803 | 74.66 | 0.8351 |
| nrdA_333/2370 | 74.22 | 0.8302 |
| dnaJ_318/1122 | 74.21 | 0.8301 |
| hisG_453/900 | 73.89 | 0.8264 |
| proS_1614/1707 | 73.59 | 0.823 |
| proS_1594/1707 | 73.58 | 0.8229 |
| proS_1364/1707 | 72.98 | 0.8163 |
| pnp_1347/2160 | 72.97 | 0.8162 |
| dnaJ_950/1122 | 72.97 | 0.8161 |
| purB_1071/1329 | 72.89 | 0.8153 |
| nadE_653/741 | 72.89 | 0.8152 |
| ppiB_288/483 | 72.56 | 0.8115 |
| ispB_118/894 | 72.35 | 0.8093 |
| purB_1304/1329 | 72.28 | 0.8085 |
| purB_1167/1329 | 72.27 | 0.8084 |
| pnp_930/2160 | 72.26 | 0.8083 |
| pheT_268/2322 | 71.34 | 0.7979 |
| pheT_218/2322 | 71.33 | 0.7978 |
| ruvC_342/555 | 71.32 | 0.7977 |
| ruvC_324/555 | 71.02 | 0.7943 |
| mdtJ_198/342 | 69.96 | 0.7825 |

| | | |
|---|---|---|
| cfiB_194/1446 | 69.95 | 0.7824 |
| pseG_341/825 | 69.95 | 0.7823 |
| ttcA_621/756 | 69.24 | 0.7744 |
| ahpC_397/597 | 68.39 | 0.765 |
| ackA_165/1191 | 68.25 | 0.7634 |
| arsB_327/1287 | 68.02 | 0.7608 |
| dnaJ_522/1122 | 68.01 | 0.7607 |
| purQ_425/672 | 67.80 | 0.7583 |
| ttcA_715/756 | 67.63 | 0.7564 |
| kpsD_498/1659 | 67.62 | 0.7563 |
| ahpC_444/597 | 67.24 | 0.752 |
| porA_541/1275 | 67.22 | 0.7518 |
| rimO_629/1320 | 66.53 | 0.7442 |
| rimO_429/1320 | 66.52 | 0.7441 |
| flaA_1_1581/1719 | 66.51 | 0.744 |
| mdtJ_280/342 | 66.42 | 0.7429 |
| nth_2_165/687 | 66.24 | 0.7409 |
| glgC_2_590/762 | 66.09 | 0.7392 |
| arsB_726/1287 | 66.02 | 0.7384 |
| arsB_661/1287 | 65.96 | 0.7378 |
| arsB_636/1287 | 65.96 | 0.7377 |
| arsB_608/1287 | 65.95 | 0.7376 |
| fumC_657/1392 | 65.75 | 0.7355 |
| nrdA_1317/2370 | 65.60 | 0.7337 |
| glgC_2_372/762 | 65.28 | 0.7302 |
| ruvC_337/555 | 65.28 | 0.7301 |
| pcaB_78/1368 | 65.11 | 0.7283 |
| kdsD_294/948 | 65.08 | 0.7279 |
| ahpC_558/597 | 64.67 | 0.7233 |
| ahpC_264/597 | 64.66 | 0.7232 |
| nrdA_1206/2370 | 64.48 | 0.7212 |
| parB_429/837 | 64.45 | 0.7208 |
| proC_652/732 | 64.44 | 0.7208 |
| parB_414/837 | 64.44 | 0.7207 |
| parB_333/837 | 64.43 | 0.7207 |
| mnmG_1578/1860 | 64.42 | 0.7206 |
| cfiB_954/1446 | 64.41 | 0.7205 |
| arsB_291/1287 | 64.40 | 0.7203 |
| arsB_137/1287 | 64.39 | 0.7202 |
| flaA_2_870/1719 | 63.95 | 0.7153 |
| ttuB_870/1233 | 63.36 | 0.7087 |
| thiG_428/777 | 63.24 | 0.7073 |
| rimO_1063/1320 | 63.24 | 0.7073 |
| thiG_272/777 | 63.23 | 0.7072 |
| nrdA_1053/2370 | 63.23 | 0.7072 |

| | | |
|---|---|---|
| nrdA_1047/2370 | 63.22 | 0.7071 |
| dksA_57/363 | 63.21 | 0.707 |
| rlmB_577/684 | 63.11 | 0.7059 |
| mnmG_1764/1860 | 63.08 | 0.7055 |
| arsB_276/1287 | 63.07 | 0.7054 |
| thiG_243/777 | 62.90 | 0.7035 |
| mnmG_1347/1860 | 62.83 | 0.7027 |
| mdtI_74/309 | 62.82 | 0.7027 |
| kdpC_363/411 | 62.81 | 0.7026 |
| kdpC_180/411 | 62.80 | 0.7025 |
| birA_402/654 | 62.80 | 0.7024 |
| kdsD_153/948 | 62.63 | 0.7005 |
| fmt_293/918 | 62.57 | 0.6999 |
| fmt_375/918 | 61.96 | 0.693 |
| lpxC_270/885 | 61.80 | 0.6912 |
| dapE_897/1098 | 61.66 | 0.6897 |
| argS_1197/1593 | 61.43 | 0.6871 |
| mnmG_1521/1860 | 61.42 | 0.687 |
| mnmG_1476/1860 | 61.42 | 0.6869 |
| mnmG_1455/1860 | 61.41 | 0.6868 |
| ackA_807/1191 | 61.40 | 0.6868 |
| mnmG_1437/1860 | 61.40 | 0.6868 |
| arsB_450/1287 | 61.39 | 0.6867 |
| ackA_825/1191 | 61.39 | 0.6866 |
| thiD_520/813 | 61.34 | 0.6861 |
| citN_821/1347 | 60.91 | 0.6813 |
| recN_540/1524 | 60.71 | 0.6791 |
| argD_625/1182 | 60.70 | 0.679 |
| parB_63/837 | 60.54 | 0.6771 |
| korA_822/1125 | 60.53 | 0.677 |
| recN_738/1524 | 60.52 | 0.6769 |
| prmC_345/816 | 60.38 | 0.6753 |
| nrdA_357/2370 | 60.37 | 0.6752 |
| htpX_974/1188 | 60.36 | 0.6751 |
| dltA_403/1509 | 60.35 | 0.675 |
| mnmG_669/1860 | 59.73 | 0.6681 |
| dapE_1065/1098 | 59.69 | 0.6677 |
| hemC_750/924 | 59.64 | 0.6671 |
| thiG_331/777 | 59.58 | 0.6664 |
| thiD_580/813 | 59.57 | 0.6663 |
| glgC_2_282/762 | 59.56 | 0.6662 |
| ackA_789/1191 | 59.50 | 0.6655 |
| arsB_796/1287 | 59.50 | 0.6655 |
| arsB_715/1287 | 59.49 | 0.6654 |
| arsB_616/1287 | 59.48 | 0.6653 |

| | | |
|---|---|---|
| rcsC_2238/2310 | 59.11 | 0.6612 |
| pleD_81/1245 | 59.10 | 0.6611 |
| yrdA_405/549 | 59.10 | 0.661 |
| pleD_585/1245 | 59.10 | 0.661 |
| cydB_373/1125 | 59.09 | 0.6609 |
| recN_51/1524 | 59.09 | 0.6609 |
| infA_80/219 | 59.09 | 0.6609 |
| copA_1_1610/2352 | 59.08 | 0.6609 |
| infA_33/219 | 59.08 | 0.6608 |
| argH_1045/1383 | 59.08 | 0.6608 |
| ftsA_501/1383 | 59.07 | 0.6607 |
| soj_435/786 | 58.97 | 0.6596 |
| soj_123/786 | 58.97 | 0.6596 |
| soj_351/786 | 58.96 | 0.6595 |
| queE_385/744 | 58.96 | 0.6595 |
| soj_325/786 | 58.96 | 0.6594 |
| porA_187/1275 | 58.95 | 0.6594 |
| birA_603/654 | 58.95 | 0.6594 |
| soj_144/786 | 58.95 | 0.6593 |
| mntB_678/804 | 58.95 | 0.6593 |
| atpC_183/390 | 58.94 | 0.6593 |
| fmt_867/918 | 58.94 | 0.6592 |
| aspS_250/1752 | 58.94 | 0.6592 |
| fabF_189/1215 | 58.93 | 0.6591 |
| aat_2116/2130 | 58.93 | 0.6591 |
| aspS_706/1752 | 58.83 | 0.658 |
| aspS_687/1752 | 58.82 | 0.6579 |
| soj_387/786 | 58.81 | 0.6578 |
| sucD_606/870 | 58.71 | 0.6567 |
| sucD_537/870 | 58.70 | 0.6566 |
| sucD_624/870 | 58.70 | 0.6565 |
| pal_20/498 | 58.69 | 0.6565 |
| sucD_613/870 | 58.69 | 0.6564 |
| mutS2_378/2208 | 58.69 | 0.6564 |
| lpxC_633/885 | 58.68 | 0.6563 |
| recN_1225/1524 | 58.58 | 0.6552 |
| dcuB_136/1425 | 58.50 | 0.6544 |
| sucD_408/870 | 58.50 | 0.6543 |
| dcuB_102/1425 | 58.50 | 0.6543 |
| rpoN_565/1251 | 58.49 | 0.6542 |
| clpB_675/2574 | 58.49 | 0.6542 |
| flaA_2_742/1719 | 58.48 | 0.6541 |
| flaA_2_723/1719 | 58.42 | 0.6534 |
| rlmB_276/684 | 58.19 | 0.6509 |
| feoB_1098/1842 | 58.18 | 0.6508 |

| | | |
|---|---|---|
| aer_1_65/498 | 58.15 | 0.6504 |
| aer_1_21/498 | 58.14 | 0.6503 |
| flaA_2_826/1719 | 57.92 | 0.6478 |
| rocA_2252/3489 | 57.72 | 0.6456 |
| uvrB_189/1974 | 57.53 | 0.6434 |
| yycB_1045/1164 | 57.52 | 0.6433 |
| uvrB_1704/1974 | 57.52 | 0.6433 |
| yajR_935/1296 | 57.51 | 0.6433 |
| uvrB_1513/1974 | 57.51 | 0.6432 |
| tolB_212/1209 | 57.51 | 0.6432 |
| uvrB_943/1974 | 57.50 | 0.6432 |
| uvrB_120/1974 | 57.50 | 0.6431 |
| tolB_132/1209 | 57.50 | 0.6431 |
| uvrB_1089/1974 | 57.49 | 0.6431 |
| sucC_48/1164 | 57.49 | 0.643 |
| rpoBC_2_3641/4554 | 57.49 | 0.643 |
| ttcA_105/756 | 57.49 | 0.643 |
| sucC_237/1164 | 57.48 | 0.6429 |
| rhaM_131/318 | 57.48 | 0.6429 |
| rpsA_110/1671 | 57.48 | 0.6429 |
| proP_4_560/1296 | 57.47 | 0.6428 |
| moaA_201/963 | 57.47 | 0.6428 |
| rpoBC_2_3850/4554 | 57.47 | 0.6428 |
| proP_2_1116/1362 | 57.47 | 0.6428 |
| mnmA_1_845/1017 | 57.46 | 0.6427 |
| plsY_460/609 | 57.46 | 0.6427 |
| mdh_633/903 | 57.46 | 0.6426 |
| kdpD_30/1821 | 57.45 | 0.6426 |
| murD_1080/1209 | 57.45 | 0.6426 |
| kdpD_702/1821 | 57.45 | 0.6426 |
| kdpD_1798/1821 | 57.45 | 0.6425 |
| kdpD_559/1821 | 57.44 | 0.6425 |
| kdpB_948/2046 | 57.44 | 0.6424 |
| icd2_390/2205 | 57.44 | 0.6424 |
| kdpD_4/1821 | 57.43 | 0.6424 |
| kdpB_1874/2046 | 57.43 | 0.6424 |
| htpX_655/1188 | 57.43 | 0.6423 |
| kdpB_1391/2046 | 57.42 | 0.6423 |
| htpX_143/1188 | 57.42 | 0.6422 |
| frdC_727/783 | 57.42 | 0.6422 |
| icd2_700/2205 | 57.42 | 0.6422 |
| hisS_1012/1227 | 57.41 | 0.6422 |
| ffh_428/1338 | 57.41 | 0.6421 |
| gpmI_506/1479 | 57.41 | 0.6421 |
| dsbL_1_78/360 | 57.40 | 0.6421 |

| | | |
|---|---|---|
| btuD_1_265/855 | 57.40 | 0.642 |
| fusA_1331/2076 | 57.40 | 0.642 |
| dnaE_3535/3603 | 57.40 | 0.642 |
| atpC_290/390 | 57.39 | 0.6419 |
| copA_1_1242/2352 | 57.39 | 0.6419 |
| atpC_253/390 | 57.39 | 0.6419 |
| clpA_1936/2130 | 57.38 | 0.6418 |
| argC_58/1029 | 57.38 | 0.6418 |
| ccpA_24/1026 | 57.38 | 0.6418 |
| atoC_1248/1302 | 57.38 | 0.6418 |
| aas_3474/3513 | 57.38 | 0.6417 |
| aroA_66/1287 | 57.37 | 0.6417 |
| aroA_36/1287 | 57.36 | 0.6416 |
| uvrB_1866/1974 | 57.20 | 0.6398 |
| murC_1290/1299 | 57.20 | 0.6397 |
| lpxC_483/885 | 57.19 | 0.6396 |
| kdpB_674/2046 | 57.19 | 0.6396 |
| kdpD_615/1821 | 57.18 | 0.6395 |
| kdpB_1818/2046 | 57.18 | 0.6395 |
| kdpD_1593/1821 | 57.17 | 0.6395 |
| kdpB_1614/2046 | 57.17 | 0.6394 |
| kdpB_1063/2046 | 57.17 | 0.6394 |
| kdpB_723/2046 | 57.16 | 0.6394 |
| kdpB_1494/2046 | 57.16 | 0.6393 |
| htpX_399/1188 | 57.16 | 0.6393 |
| kdpB_1453/2046 | 57.15 | 0.6392 |
| kdpB_1139/2046 | 57.14 | 0.6392 |
| tmk_380/579 | 57.10 | 0.6386 |
| dcuB_24/1425 | 57.09 | 0.6385 |
| pfs_528/687 | 57.04 | 0.638 |
| kdpA_1_179/261 | 57.03 | 0.6379 |
| eptA_1119/1539 | 56.94 | 0.6369 |
| cfiB_237/1446 | 56.90 | 0.6364 |
| thiG_474/777 | 56.86 | 0.636 |
| mnmA_1_426/1017 | 56.86 | 0.636 |
| flaA_2_855/1719 | 56.45 | 0.6313 |
| flaA_2_819/1719 | 56.44 | 0.6313 |
| aer_1_234/498 | 56.43 | 0.6312 |
| ackA_456/1191 | 56.09 | 0.6273 |
| yafP_79/459 | 55.94 | 0.6257 |
| flaA_2_1554/1719 | 55.93 | 0.6256 |
| dapA_1_162/909 | 55.92 | 0.6255 |
| atpD_1_437/522 | 55.92 | 0.6254 |
| thiG_171/777 | 55.86 | 0.6248 |
| smpB_189/453 | 55.85 | 0.6247 |

| | | |
|---|---|---|
| yflS_134/168 | 55.85 | 0.6246 |
| rpoBC_1_4034/4128 | 55.84 | 0.6246 |
| gppA_2_197/975 | 55.84 | 0.6246 |
| tsf_246/1074 | 55.84 | 0.6246 |
| kgtP_424/1260 | 55.84 | 0.6245 |
| glyS_1573/1995 | 55.84 | 0.6245 |
| hom_842/1248 | 55.83 | 0.6244 |
| glmU_27/1290 | 55.83 | 0.6244 |
| aas_2194/3513 | 55.83 | 0.6244 |
| hom_589/1248 | 55.82 | 0.6244 |
| dapE_717/1098 | 55.82 | 0.6243 |
| aas_1587/3513 | 55.82 | 0.6243 |
| carB_1236/3270 | 55.81 | 0.6243 |
| bioB_183/837 | 55.80 | 0.6242 |
| mog_477/543 | 55.78 | 0.6239 |
| copA_1_837/2352 | 55.78 | 0.6239 |
| copA_1_783/2352 | 55.77 | 0.6238 |
| bamA_294/2220 | 55.77 | 0.6237 |
| copA_1_1335/2352 | 55.76 | 0.6237 |
| bamA_279/2220 | 55.76 | 0.6237 |
| clpA_1341/2130 | 55.75 | 0.6236 |
| atpC_102/390 | 55.75 | 0.6236 |
| birA_348/654 | 55.74 | 0.6235 |
| asd_867/1032 | 55.74 | 0.6235 |

| | relative importance | scaled importance |
|---|---|---|
| porA_513/1275 | 89.3906 | 0.9998 |
| porA_803/1275 | 86.0156 | 0.9621 |
| porA_592/1275 | 84.5313 | 0.9455 |
| porA_548/1275 | 83.5625 | 0.9346 |
| porA_445/1275 | 83.1875 | 0.9304 |
| porA_562/1275 | 82.1992 | 0.9194 |
| porA_580/1275 | 81.2031 | 0.9082 |
| porA_400/1275 | 81.1563 | 0.9077 |
| porA_250/1275 | 79.457 | 0.8887 |
| porA_459/1275 | 79.2383 | 0.8863 |
| porA_411/1275 | 75.9004 | 0.8489 |
| porA_541/1275 | 67.2172 | 0.7518 |
| porA_187/1275 | 58.9543 | 0.6594 |

| percentage |
| --- |
| 0.0048 |
| 0.0046 |
| 0.0045 |
| 0.0044 |
| 0.0044 |
| 0.0044 |
| 0.0043 |
| 0.0043 |
| 0.0042 |
| 0.0042 |
| 0.004 |
| 0.0036 |
| 0.0031 |

XGBOOST PARAMETERS

| *name* | *value* |
|---|---|
| silent | TRUE |
| eta | 0.3 |
| colsample_bylevel | 1 |
| objective | binary:logistic |
| min_child_weight | 1 |
| nthread | 8 |
| seed | 27234256 |
| max_depth | 6 |
| colsample_bytree | 1 |
| lambda | 1 |
| gamma | 0 |
| alpha | 0 |
| booster | gbtree |
| grow_policy | depthwise |
| nround | 50 |
| subsample | 1 |
| max_delta_step | 0 |
