## Supplementary material for "Biological machine learning combined with bacterial population genomics reveals common and rare allelic variants of genes to cause disease": supp table 2

| Strain Name | Host | Disease | Source | Region | Year | Accession |
|---|---|---|---|---|---|---|
| CA6e | Sheep | Abortion | Aborted placenta | CA,USA | 1991 | SRR3094400 |
| CA7e | Sheep | Abortion | Aborted placenta | CA,USA | 1992 | SRR3094401 |
| ID1 | Sheep | Abortion | Aborted placenta | ID,USA | 1992 | SRR3094402 |
| CA8e | Sheep | Abortion | Aborted placenta | CA,USA | 1993 | SRR3094403 |
| ID12 | Sheep | Abortion | Aborted placenta | ID,USA | 1993 | SRR3094404 |
| ID15 | Sheep | Abortion | Aborted placenta | ID,USA | 1993 | SRR3094405 |
| ID28 | Sheep | Abortion | Aborted placenta | ID,USA | 1993 | SRR3094406 |
| ID31 | Sheep | Abortion | Aborted placenta | ID,USA | 1993 | SRR3094407 |
| ID34 | Sheep | Abortion | Aborted placenta | ID,USA | 1993 | SRR3094408 |
| CA3e | Sheep | Abortion | Aborted placenta | CA,USA | 1999 | SRR3094409 |
| CA4e | Sheep | Abortion | Aborted placenta | CA,USA | 1999 | SRR3094410 |
| CA5e | Sheep | Abortion | Aborted placenta | CA,USA | 2000 | SRR3094411 |
| VDL705 | Sheep | Abortion | Aborted placenta | IA,USA | 2003 | SRR3094412 |
| VLD698 | Sheep | Abortion | Aborted placenta | IA,USA | 2003 | SRR3094413 |
| CA7 | Sheep | Abortion | Aborted placenta | CA,USA | 2003 | SRR3094414 |
| ID181077-B | Sheep | Abortion | Aborted placenta | ID,USA | 2004 | SRR3094415 |
| ID323147-F | Sheep | Abortion | Aborted placenta | ID,USA | 2004 | SRR3094416 |
| VDL5908 | Sheep | Abortion | Aborted placenta | IA,USA | 2004 | SRR3094417 |
| VDL3842 | Sheep | Abortion | Aborted placenta | IA,USA | 2004 | SRR3094418 |
| CA5 | Sheep | Abortion | Aborted placenta | CA,USA | 2004 | SRR3094419 |
| CA8 | Sheep | Abortion | Aborted placenta | CA,USA | 2004 | SRR3094420 |
| ID219083-A | Sheep | Abortion | Aborted placenta | ID,USA | 2005 | SRR3094421 |
| VDL3080 | Sheep | Abortion | Aborted placenta | IA,USA | 2005 | SRR3094422 |
| SD3831 | Sheep | Abortion | Aborted placenta | SD,USA | 2005 | SRR3094423 |
| CA3 | Sheep | Abortion | Aborted placenta | CA,USA | 2005 | SRR3094424 |
| ID273138-G | Sheep | Abortion | Aborted placenta | ID,USA | 2006 | SRR3094425 |
| ID428211-A | Sheep | Abortion | Aborted placenta | ID,USA | 2006 | SRR3094426 |
| ID147052-A | Sheep | Abortion | Aborted placenta | ID,USA | 2007 | SRR3094427 |
| VDL6220 | Sheep | Abortion | Aborted placenta | IA,USA | 2007 | SRR3094428 |
| ND6 | Sheep | Abortion | Aborted placenta | ND,USA | 2007 | SRR3094429 |
| VDL2192 | Sheep | Abortion | Aborted placenta | IA,USA | 2008 | SRR3094430 |
| VDL8958 | Sheep | Abortion | Aborted placenta | IA,USA | 2009 | SRR3094431 |
| VDL35044 | Sheep | Abortion | Aborted placenta | IA,USA | 2009 | SRR3094432 |
| VDL1957 | Sheep | Abortion | Aborted placenta | IA,USA | 2010 | SRR3094433 |
| VDL2764 | Sheep | Abortion | Aborted placenta | IA,USA | 2010 | SRR3094434 |
| VDL5414 | Sheep | Abortion | Aborted placenta | IA,USA | 2011 | SRR3094435 |
| CO351 | Sheep | Abortion | Aborted placenta | CO,USA | 2011 | SRR3094436 |
| ID33 | Sheep | Abortion | Aborted placenta | ID,USA | 1993 | SRR3094437 |
| ID8 | Sheep | Abortion | Aborted placenta | ID,USA | 1993 | SRR3094438 |
| ID30 | Sheep | Abortion | Aborted placenta | ID,USA | 1993 | SRR3094439 |
| ID10 | Sheep | Abortion | Aborted placenta | ID,USA | 1993 | SRR3094440 |
| CA1e | Sheep | Abortion | Aborted placenta | CA,USA | 1998 | SRR3094441 |
| CA2e | Sheep | Abortion | Aborted placenta | CA,USA | 1998 | SRR3094442 |
| ID017002-A | Sheep | Abortion | Aborted placenta | ID,USA | 2004 | SRR3094443 |
| CA2 | Sheep | Abortion | Aborted placenta | CA,USA | 2003 | SRR3094444 |
| VDL4646 | Sheep | Abortion | Aborted placenta | IA,USA | 2004 | SRR3094445 |
| VDL2401 | Sheep | Abortion | Aborted placenta | IA,USA | 2007 | SRR3094446 |
| ND7 | Sheep | Abortion | Aborted placenta | ND,USA | 2007 | SRR3094447 |
| ND9 | Sheep | Abortion | Aborted placenta | ND,USA | 2008 | SRR3094448 |
| VDL902 | Sheep | Abortion | Aborted placenta | IA,USA | 2008 | SRR3094449 |
| VDL2019 | Sheep | Abortion | Aborted placenta | IA,USA | 2008 | SRR3094450 |

| | | | | | | |
|---|---|---|---|---|---|---|
| VDL213 | Sheep | Abortion | Aborted placenta | IA, USA | 2009 | SRR3094451 |
| VDL2945 | Sheep | Abortion | Aborted placenta | IA, USA | 2009 | SRR3094452 |
| VDL6069 | Sheep | Abortion | Aborted placenta | IA, USA | 2010 | SRR3094453 |
| CA10 | Cattle | Abortion | Aborted placenta | CA, USA | 2003 | SRR3094454 |
| CA12 | Cattle | Abortion | Aborted placenta | CA, USA | 2003 | SRR3094455 |
| CA11 | Cattle | Abortion | Aborted placenta | CA, USA | 2004 | SRR3094456 |
| ND3 | Cattle | Abortion | Aborted placenta | ND, USA | 2005 | SRR3094457 |
| VDL31248 | Cattle | Abortion | Aborted placenta | IA, USA | 2009 | SRR3094458 |
| VDL2738 | Goat | Abortion | Aborted placenta | IA, USA | 2005 | SRR3094459 |
| VDL1625 | Goat | Abortion | Aborted placenta | IA, USA | 2008 | SRR3094460 |
| VDL4023 | Goat | Abortion | Aborted placenta | IA, USA | 2010 | SRR3094461 |
| VDL4350 | Goat | Abortion | Aborted placenta | IA, USA | 2010 | SRR3094462 |
| UK33 | Sheep | Abortion | Aborted placenta | UK | 2006 | SRR3094476 |
| UK29 | Sheep | Abortion | Aborted placenta | UK | 2005 | SRR3094477 |
| UK19 | Sheep | Abortion | Aborted placenta | UK | 2004 | SRR3094478 |
| VDL35490 | Sheep | Abortion | Aborted placenta | IA, USA | 2012 | SRR3094480 |
| VDL3452 | Sheep | Abortion | Aborted placenta | IA, USA | 2013 | SRR3094481 |
| VDL6794 | Sheep | Abortion | Aborted placenta | IA, USA | 2013 | SRR3094482 |
| UK40 | Sheep | Abortion | Aborted placenta | UK | 2008 | SRR3094483 |
| UK11 | Sheep | Abortion | Aborted placenta | UK | 2003 | SRR3094484 |
| UK10 | Sheep | Abortion | Aborted placenta | UK | 2003 | SRR3094485 |
| UK24 | Sheep | Abortion | Aborted placenta | UK | 2005 | SRR3094486 |
| UK32 | Sheep | Abortion | Aborted placenta | UK | 2006 | SRR3094487 |
| UK37 | Sheep | Abortion | Aborted placenta | UK | 2007 | SRR3094488 |
| ID7 | Sheep | Abortion | Aborted placenta | ID, USA | 1992 | SRR3094489 |
| ID27 | Sheep | Abortion | Aborted placenta | ID, USA | 1993 | SRR3094490 |
| ID25 | Sheep | Abortion | Aborted placenta | ID, USA | 1993 | SRR3094491 |
| SD4165 | Sheep | Abortion | Aborted placenta | SD, USA | 2005 | SRR3094492 |
| VDL2847 | Sheep | Abortion | Aborted placenta | IA, USA | 2010 | SRR3094493 |
| VDL2918 | Sheep | Abortion | Aborted placenta | IA, USA | 2005 | SRR3094494 |
| 302 | Sheep | Abortion | Aborted placenta | IA, USA | 2014 | SRR3094496 |
| 3008 | Sheep | Abortion | Aborted placenta | IA, USA | 2014 | SRR3094497 |
| 5406 | Sheep | Abortion | Aborted placenta | MI, USA | 2014 | SRR3094498 |
