## Supplementary material for "Biological machine learning combined with bacterial population genomics reveals common and rare allelic variants of genes to cause disease": Supp table 3

| Genome | Collection D | Geographic Location |
|---|---|---|
| 197.14907 | 2018 | USA:PA |
| 197.15041 | 2018 | USA:MN |
| 197.15927 | 2016 | USA:TX |
| 197.15936 | 2016 | USA:PA |
| 197.15943 | 2016 | USA:TN |
| 197.15952 | 2016 | USA:PA |
| 197.15957 | 2016 | USA:OR |
| 197.15958 | 2016 | USA:NE |
| 197.15964 | 2016 | USA:ID |
| 197.15966 | 2016 | USA:CO |
| 197.15972 | 2016 | USA:NJ |
| 197.15982 | 2016 | USA:TX |
| 197.15983 | 2016 | USA:PA |
| 197.15988 | 2016 | USA:MI |
| 197.15993 | 2016 | USA:PA |
| 197.15994 | 2016 | USA:MI |
| 197.15997 | 2016 | USA:NY |
| 197.15999 | 2016 | USA:OH |
| 197.16001 | 2016 | USA:GA |
| 197.16005 | 2016 | USA:OH |
| 197.16021 | 2016 | USA:PA |
| 197.16023 | 2016 | USA:PA |
| 197.16031 | 2016 | USA:WI |
| 197.16033 | 2016 | USA:CA |
| 197.16041 | 2016 | USA:TX |
| 197.16042 | 2016 | USA:NE |
| 197.16048 | 2016 | USA:TX |
| 197.16051 | 2016 | USA:WA |
| 197.16053 | 2016 | USA:MI |
| 197.16058 | 2016 | USA:NJ |
| 197.16060 | 2016 | USA:WA |
| 197.16064 | 2016 | USA:NE |
| 197.16072 | 2016 | USA:MI |
| 197.16074 | 2016 | USA:ID |
| 197.16075 | 2016 | USA:MI |
| 197.16078 | 2016 | USA:CA |
| 197.16079 | 2016 | USA:OR |
| 197.16084 | 2016 | USA:FL |
| 197.16099 | 2016 | USA:NJ |
| 197.16103 | 2016 | USA:TX |
| 197.16118 | 2016 | USA:WI |
| 197.16127 | 2016 | USA:MN |

| | | |
|---|---|---|
| 197.16139 | 2016 | USA:WI |
| 197.16141 | 2016 | USA:UT |
| 197.16145 | 2016 | USA:AZ |
| 197.16146 | 2016 | USA:CA |
| 197.16155 | 2016 | USA:TX |
| 197.16156 | 2016 | USA:OH |
| 197.16157 | 2016 | USA:WA |
| 197.16159 | 2016 | USA:MI |
| 197.16162 | 2016 | USA:UT |
| 197.16165 | 2016 | USA:MD |
| 197.16169 | 2016 | USA:TX |
| 197.16171 | 2016 | USA:GA |
| 197.16180 | 2016 | USA:NE |
| 197.16183 | 2016 | USA:WA |
| 197.16185 | 2016 | USA:SD |
| 197.16192 | 2016 | USA:TX |
| 197.16199 | 2016 | USA:OR |
| 197.16209 | 2016 | USA:WA |
| 197.16210 | 2016 | USA:SD |
| 197.16211 | 2016 | USA:TX |
| 197.16215 | 2016 | USA:WI |
| 197.16221 | 2016 | USA:CA |
| 197.16223 | 2016 | USA:WA |
| 197.16225 | 2016 | USA:MD |
| 197.16227 | 2016 | USA:MN |
| 197.16230 | 2016 | USA:PA |
| 197.16231 | 2016 | USA:CA |
| 197.16233 | 2016 | USA:ID |
| 197.16237 | 2016 | USA:SC |
| 197.16239 | 2016 | USA:OR |
| 197.16242 | 2016 | USA:WI |
| 197.16248 | 2016 | USA:PA |
| 197.16251 | 2016 | USA:CA |
| 197.16259 | 2016 | USA:NY |
| 197.16263 | 2016 | USA:WA |
| 197.16268 | 2016 | USA:PA |
| 197.16269 | 2016 | USA:CO |
| 197.16275 | 2016 | USA:WI |
| 197.16280 | 2016 | USA:WI |
| 197.16287 | 2016 | USA:CO |
| 197.16291 | 2016 | USA:ID |
| 197.16300 | 2016 | USA:TX |
| 197.16303 | 2016 | USA:PA |

| | | |
|---|---|---|
| 197.16307 | 2016 | USA:TX |
| 197.16310 | 2016 | USA:SD |
| 197.16313 | 2016 | USA:AZ |
| 197.16322 | 2016 | USA:TN |
| 197.16334 | 2016 | USA:WA |
| 197.16341 | 2016 | USA:CO |
| 197.16350 | 2016 | USA:MN |
| 197.16351 | 2016 | USA:MI |
| 197.16353 | 2016 | USA:WI |
| 197.16358 | 2016 | USA:AZ |
| 197.16362 | 2016 | USA:CA |
| 197.16364 | 2016 | USA:OR |
| 197.16365 | 2016 | USA:NJ |
| 197.16366 | 2016 | USA:ID |
| 197.16372 | 2016 | USA:NE |
| 197.16373 | 2016 | USA:PA |
| 197.16374 | 2016 | USA:ID |
| 197.16380 | 2016 | USA:TX |
| 197.16387 | 2016 | USA:TX |
| 197.16389 | 2016 | USA:NY |
| 197.16390 | 2016 | USA:PA |
| 197.16392 | 2016 | USA:PA |

197.14907
197.15041
197.15927
197.15936
197.15943
197.15952
197.15957
197.15958
197.15964
197.15966
197.15972
197.15982
197.15983
197.15988
197.15993
197.15994
197.15997
197.15999
197.16001
197.16005
197.16021
197.16023
197.16031
197.16033
197.16041
197.16042
197.16048
197.16051
197.16053
197.16058
197.16060
197.16064
197.16072
197.16074
197.16075
197.16078
197.16079
197.16084
197.16099
197.16103
197.16118
197.16127
197.16139
197.16141
197.16145

197.16146
197.16155
197.16156
197.16157
197.16159
197.16162
197.16165
197.16169
197.16171
197.16180
197.16183
197.16185
197.16192
197.16199
197.16209
197.16210
197.16211
197.16215
197.16221
197.16223
197.16225
197.16227
197.16230
197.16231
197.16233
197.16237
197.16239
197.16242
197.16248
197.16251
197.16259
197.16263
197.16268
197.16269
197.16275
197.16280
197.16287
197.16291
197.16300
197.16303
197.16307
197.16310
197.16313
197.16322
197.16334

197.16341
197.16350
197.16351
197.16353
197.16358
197.16362
197.16364
197.16365
197.16366
197.16372
197.16373
197.16374
197.16380
197.16387
197.16389
197.16390
197.16392
