## Supplementary material for "Biological machine learning combined with bacterial population genomics reveals common and rare allelic variants of genes to cause disease": Supp table 4

| | Extraintestinal | Intestinal |
|---|---|---|
| **Extraintestinal** | **18358** | **389964** |
| **Intestinal** | **6003** | **821898** |

| Measure | Value | Formula |
|---|---|---|
| Sensitivity | 0.7536 | TP / (TP + FN) |
| Specificity | 0.6782 | TN / (FP + TN) |
| Precision | 0.045 | TP / (TP + FP) |
| Negative Predictive Value | 0.9927 | TN / (TN + FN) |
| False Positive Rate | 0.3218 | FP / (FP + TN) |
| False Discovery Rate | 0.955 | FP / (FP + TP) |
| False Negative Rate | 0.2464 | FN / (FN + TP) |
| Accuracy | 0.6797 | (TP + TN) / (P + N) |
| F1 Score | 0.0849 | 2TP / (2TP + FP + FN) |
| Matthews Correlation Coefficient | 0.1276 | TP*TN - FP*FN / sqrt((TP+F |

FP)*(TP+FN)*(TN+FP)*(TN+FN))

Sensitivity
Specificity
Precision
Negative Predictive Value
False Positive Rate
False Discovery Rate
False Negative Rate
Accuracy
F1 Score
Matthews Correlation Coefficient
